# The Effect of Visual Uncertainty on Implicit Motor Adaptation

**DOI:** 10.1101/2020.03.15.992008

**Authors:** Jonathan S. Tsay, Guy Avraham, Hyosub E. Kim, Darius E. Parvin, Zixuan Wang, Richard B. Ivry

## Abstract

Sensorimotor adaptation is driven by sensory prediction errors, the difference between the predicted and actual feedback. When the position of the feedback is made uncertain, adaptation is attenuated. This effect, in the context of optimal sensory integration models, has been attributed to a weakening of the error signal driving adaptation. Here we consider an alternative hypothesis, namely that uncertainty alters the perceived location of the feedback. We present two visuomotor adaptation experiments to compare these hypotheses, varying the size and uncertainty of a visual error signal. Uncertainty attenuated learning when the error size was small but had no effect when the error size was large. This pattern of results favors the hypothesis that uncertainty does not impact the strength of the error signal, but rather, leads to mis-localization of the error. We formalize these ideas to offer a novel perspective on the effect of visual uncertainty on implicit sensorimotor adaptation.

**SIGNIFICANCE STATEMENT:** Current models of sensorimotor adaptation assume that the rate of learning will be related to properties of the error signal (e.g., size, consistency, relevance). Recent evidence has challenged this view, pointing to a rigid, modular system, one that automatically recalibrates the sensorimotor map in response to movement errors, with minimal constraint. In light of these developments, this study revisits the influence of feedback uncertainty on sensorimotor adaptation. Adaptation was attenuated in response to a noisy feedback signal, but the effect was only manifest for small errors and not for large errors. This interaction suggests that uncertainty does not weaken the error signal. Rather, it may influence the perceived location of the feedback and thus the change in the sensorimotor map induced by that error. These ideas are formalized to show how the motor system remains exquisitely calibrated, even if adaptation is largely insensitive to the statistics of error signals.

## INTRODUCTION

The sensorimotor system learns to perform a wide array of goal-directed actions in the face of environmental and physiological changes. An integral part of this process takes place in an implicit manner, driven by sensory prediction error (SPE), the difference between the predicted and actual sensory feedback (Shadmehr, Smith, & Krakauer, 2010). The SPE is used to adjust an internal model of the body’s interactions with its environment, improving subsequent movements, and ensuring that the predicted consequences of these movements are better aligned with the actual feedback

Many studies have examined how variation in the error signal influences adaptation. According to an optimal integration model (OI model), the error is estimated using a weighted signal composed of visual and proprioceptive feedback (Burge, Ernst, & Banks, 2008; Körding & Wolpert, 2004; Wei & Körding, 2010). The weights in this model reflect the degree of uncertainty associated with each source (Ernst & Banks, 2002). Consider a visuomotor adaptation experiment in which the direction of visual feedback is rotated relative to the direction of the hand, resulting in a mismatch (error signal) between the perturbed visual feedback and veridical proprioceptive feedback. The relative reliability assigned to each input determines the perceived position of the hand. For example, a noisy visual signal will reduce the weight given to the visual feedback, biasing the estimate towards proprioceptive feedback. Based on the difference between this estimate and the predicted feedback location (i.e., the target), the motor system corrects for a constant proportion of the error, producing a linear relationship between the amount of trial-to-trial adaptation and the error size (Fig. 1a). In the simplest form of the OI model, the motor system learns from all error sizes in the same way (i.e., constant learning rate) such that the sensorimotor map updates proportionally as a function of error size (Cheng & Sabes, 2006; Tanaka, Krakauer, & Sejnowski, 2012; Thoroughman & Shadmehr, 2000). When the perturbed visual feedback is made more uncertain, the visual error signal is weakened and consequently, learning is reduced. The impact of uncertainty will be evident for all error sizes (Fig. 1a) producing an overall attenuation of the learning function.

**Fig. 1.**
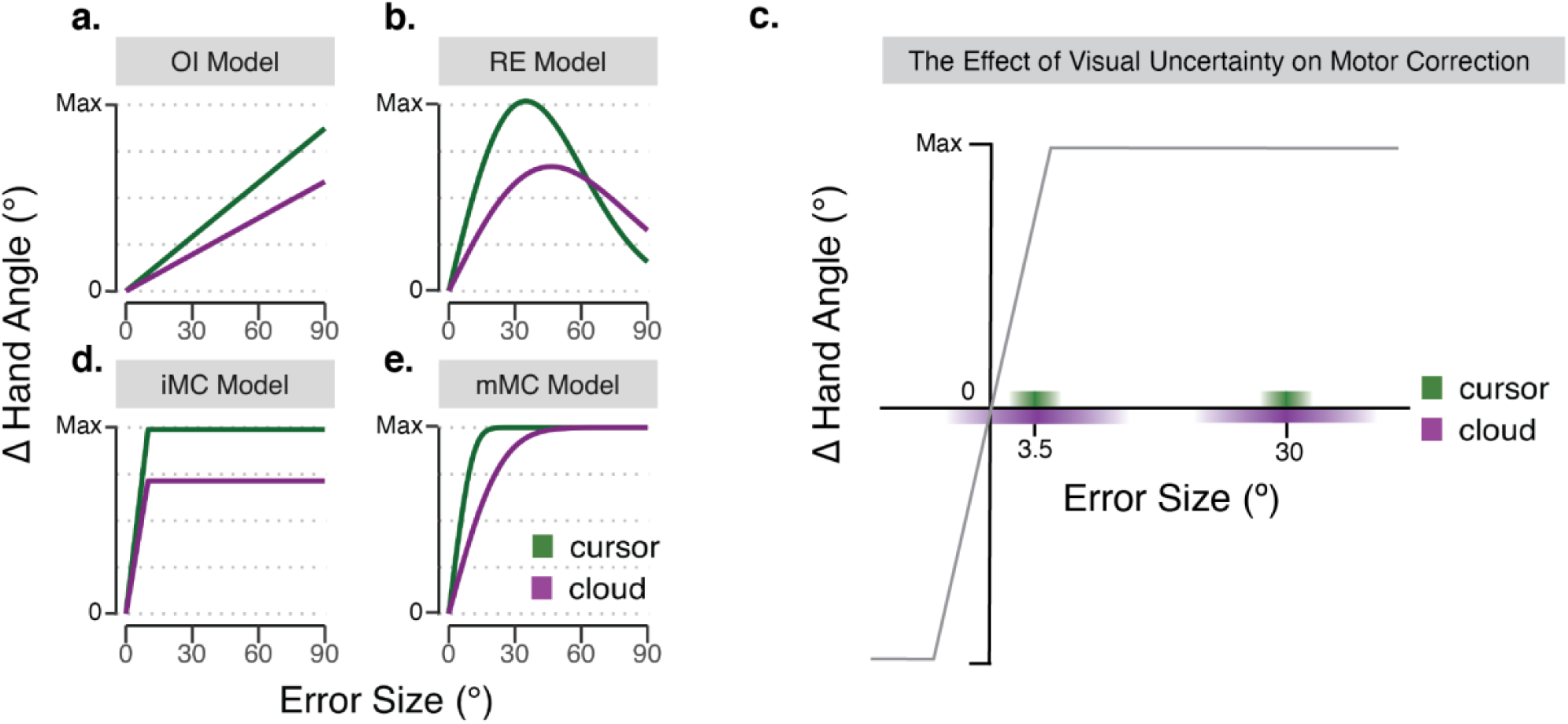
Modeling the effect of visual uncertainty on visuomotor adaptation. Panels a-d display the predicted trial-by-trial change in hand angle as a function of error size and visual uncertainty (cursor = certain feedback, cloud = uncertain feedback). The illustrative motor responses are generated by simulations of the **(a)** optimal integration model (OI model), **(b)** relevance estimation model (RE model), **(d)** integrated motor correction model (iMC model), and **(e)** the mis-localized motor correction model (mMC model). Both variants of the motor correction model **(c)** assume that the update function is composed of a linear zone, where motor updates are proportional to the size of the error, and a saturation zone, estimated to start around 5°, over which the size of the motor update is invariant. The saturation level of the colored bars on the x-axis depict the distribution of perceived locations for the low uncertainty (green) and high uncertainty (purple) conditions. For a large 30° error, the perceived location of the error always falls in the saturation zone and thus adaptation is similar for the cursor and cloud conditions. For the small 3.5°, uncertainty will impact the size of the update, including sign flips when the perceived location of the error is of the opposite sign as the actual error.

However, several studies have revealed a nonlinear relationship between the rate of adaptation and the size of the error, posing a challenge to the notion of a constant learning rate (Fine & Thoroughman, 2007; Robinson, Noto, & Bevans, 2003; Wei & Körding, 2009). One account for this nonlinearity centers on the hypothesis that the error signal reflects not only optimal multisensory integration but also a form of causal interference (Relevance Estimation Model, Fig. 1b). The RE model posits that the motor system determines whether the error is intrinsic to its performance or may be attributed to extrinsic factors, with the former more relevant for keeping the system properly calibrated. Error size provides one heuristic: Large errors are likely to be attributed to extrinsic sources (e.g., an unexpected gust of wind), especially when their occurrence is relatively low, and thus discounted. In contrast, small errors are generally attributed to noise within the motor system and thus given more weight in driving learning. Feedback uncertainty offers another heuristic: As the feedback signal becomes noisier, the source of the error signal becomes less apparent, causing the system to conservatively attribute a wider range of errors to intrinsic motor noise. For example, an increase in visual uncertainty not only reduces the weight given to the visual input, but also reduces the amount of discounting imposed on large errors. This predicts a cross-over point where the response of the motor system to large errors becomes greater to the noisy feedback signal (Fig. 1b).

Our recent studies have led an alternative computational perspective, what we will refer to as the Motor Correction Model. This model was inspired by observations that variation in the rate of adaptation is only observed over small error sizes; for errors larger than approximately 5° and ranging up to as large as 95°, the rate is essentially invariant (Kim, Morehead, Parvin, Moazzezi, & Ivry, 2018; Morehead, Taylor, Parvin, & Ivry, 2017). To account for this non-linear function, we posited that the size of the error directly modulates the trial-to-trial change in motor output (Fig. 1c). Motor responses scale for small errors, but soon reach a saturation point, the maximal update endowed to the sensorimotor map on a single trial. The invariant response to large errors is incompatible with the core predictions of the OI model (i.e., linear dependency between motor response and error size) as well as the RE model (i.e., motor response will approach zero well before 95°).

Unlike the OI and RE models, the MC model is agnostic on the issue of feedback uncertainty. There are at least two ways in which the MC model could be modified: First, in line with models of optimal integration, visual uncertainty may attenuate the corrective output; the response to an error will be lowered given the system’s reduced confidence in the validity of an error signal (iMC model: integrated motor correction model, Fig. 1d). Alternatively, visual uncertainty may not influence the efficacy of the error signal, but rather, alter the perceived location of the feedback. By this model (mMC model: mis-localized motor correction, Fig. 1e), adaptation would be unaffected by uncertainty for large errors since the perceived locations would largely fall within the saturation zone. In contrast, uncertainty would impact the response to small errors, shifting the perceived error location and, thus, the associated update value (Fig. 1c). Indeed, on some trials, the sign of the motor correction, will be reversed due to mis-localization of the error to the opposite side of the target.

Here we report two experiments designed to evaluate the impact of visual uncertainty on sensorimotor adaptation. Unlike standard visuomotor adaptation tasks, we used non-contingent, “clamped” visual feedback in which the position of the feedback cursor was independent of the participant’s movement. Participants were fully informed of this error clamp manipulation and instructed to ignore the feedback. However, the angular offset is nonetheless treated as a SPE by the motor system, resulting in implicit sensorimotor adaptation. By manipulating the size of the angular offset and uncertainty of the feedback, we can evaluate the four models outlined in Fig. 1.

## METHODS

### Participants

A total of 120 participants (52 females, mean age = 20.3 ± 2.1 years) were recruited for two experiments. The sample sizes were based on previous studies using a similar reaching task with non-contingent visual feedback (Kim et al., 2018; Morehead et al., 2017; Parvin, McDougle, Taylor, & Ivry, 2018). All participants were right-handed, as verified with the Edinburgh Handedness Inventory (Oldfield, 1971), and received course credit or financial compensation for their participation. The experimental protocol was approved by the Institutional Review Board at the University of California, Berkeley.

### Reaching Task

The participant was seated at a custom-made table that housed a horizontally mounted LCD screen (53.2 cm by 30 cm, ASUS), positioned 27 cm above a digitizing tablet (49.3 cm by 32.7 cm, Intuos 4XL; Wacom, Vancouver, WA) (Fig. 2a). Stimuli were projected onto the LCD screen. The experimental software was custom written in Matlab, using the Psychtoolbox extensions (Brainard, 1997).

**Fig. 2.**
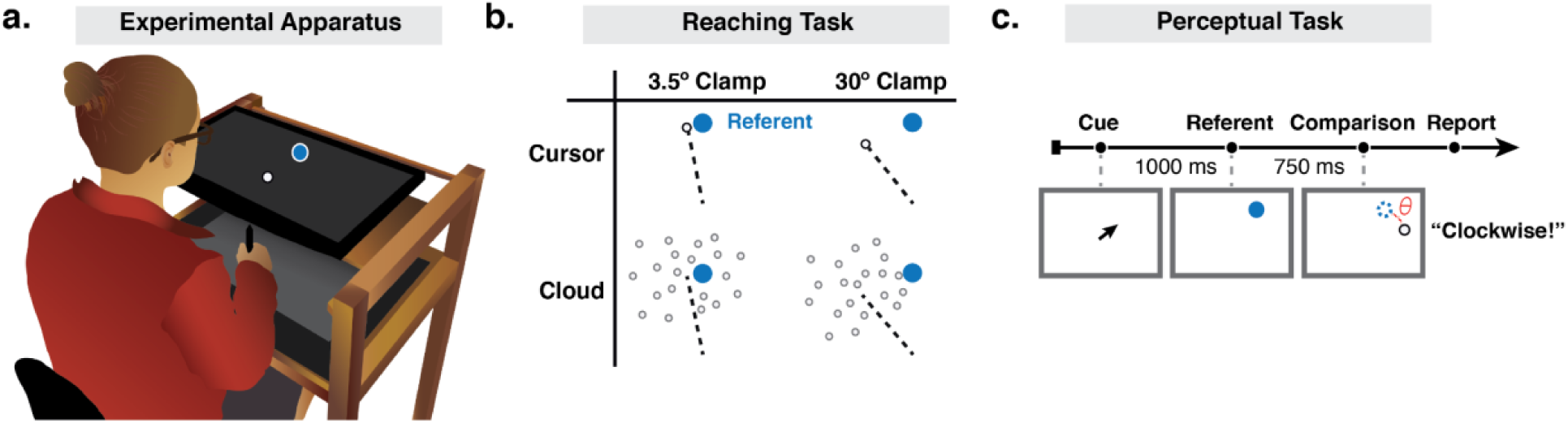
Experiment Methods. **(a)** Experimental apparatus and setup. **(b)** Schematic overview of the 2 × 2 design in Experiment 1. The referent (blue dot) was blanked from the screen before participants initiated their reach. Feedback was only presented at the endpoint, in the form of a cursor (white dot) or cloud of dots. The dotted line is included to graphically highlight the two clamp angles, pointing from the start position to the centroid of the feedback. **(c)** Trial sequence for the visual discrimination task. The centroid of the comparison stimulus (cursor or cloud) was positioned clockwise or counterclockwise relative to the target location at an angle (*θ*) equal to one of five rotation angles (see text). In the example shown, the comparison stimulus is a cursor (white circle) shifted 5° clockwise from the referent stimulus (position depicted in blue outline but not visible on the screen). The participant reported whether they perceived the centroid to be clockwise or counterclockwise relative to the remembered target location.

The participant performed center-out planar reaching movements by sliding a modified air hockey “paddle” containing an embedded stylus. The tablet recorded the position of the stylus at 200 Hz. The monitor occluded direct vision of the hand, and the room lights were extinguished to minimize peripheral vision of the arm. The participant was asked to move from a start location at the center of the workspace (indicated by a white annulus, 0.6 cm diameter) to a visual target (blue circle, 0.6 cm diameter). The target could appear at one of four locations around a virtual circle (45°, 135°, 225°, 335°), with a radial distance of 8 cm from the start location.

At the beginning of each trial, the white annulus was present on the screen, indicating the start location. The participant was instructed to move the stylus to the start location. Feedback of hand position was indicated by a solid white cursor (diameter 0.3 cm), only visible when the hand was within 2 cm of the start location. Once the start location was maintained for 500 ms, the white cursor corresponding to the participant’s hand position was turned off. The blue target appeared for only 250 ms and then blanked prior to the onset of movement. The participant was instructed to slide the stylus, attempting to rapidly “slice” through the position at which the target had appeared. We opted to blank the target prior to movement to prevent the target from being a visual reference point for the position of endpoint feedback (see also Burge et al. 2008). To ensure rapid movements, the auditory message, “too slow” was played if the movement time exceeded 300 ms. No inter-trial interval was imposed.

### Experimental Feedback Conditions

Visual feedback was limited to endpoint feedback and remained visible for 500 ms. The feedback was presented when the movement amplitude exceeded 8 cm. The radial position of the feedback was always at the distance of the target (8 cm). The angular position of the feedback was either veridical (hand position at 8 cm amplitude) or displaced from the target by a pre-specified angle (clamped perturbation). The form of the feedback was either a single white cursor (low uncertainty) or a cloud of dots (high uncertainty) (Fig. 2b). For the latter, the feedback signal was composed of a cloud of dots (25 grey 0.3 cm diameter circles) with the position of each dot pseudo-randomly drawn from a 2D isotropic Gaussian distribution with a standard deviation of 10° and with a minimal distance of 0.3 cm between dots (i.e. dots do not overlap). The center of mass of the cloud was controlled to be at the desired clamp angle on perturbation trials. The luminance of the 25 dots was adjusted such that their sum was equal to the luminance of the cursor (Körding & Wolpert, 2004; Tassinari, Hudson, & Landy, 2006; Trommershäuser, Maloney, & Landy, 2003). In addition to trials with veridical and clamped feedback, there were also trials with no feedback.

### Experiment 1

Experiment 1 (four groups, n = 24/group) was designed to examine the impact of visual uncertainty on sensorimotor adaptation when the error signal remained invariant for the duration of the experiment. A 2 × 2 factorial design was employed, with one factor based on error size (3.5° or 30° clamped displacement of feedback relative to target) and the other based on certainty (cursor or cloud). Participants were randomly assigned to one of the four conditions, and within each group, the direction of the rotation (clockwise or counterclockwise relative to the target) was counterbalanced. By using a fixed perturbation for each participant for the duration of the experiment, we could observe the learning function to near asymptotic performance.

The experiment consisted of 700 trials, divided into 5 blocks: No feedback baseline (20 trials), veridical feedback baseline (60 trials), clamped feedback perturbation (600 trials), no feedback post-perturbation (12 trials), and veridical feedback post-perturbation (8 trials). The initial baseline trials were included to familiarize the participants with the apparatus and provide veridical feedback to minimize idiosyncratic directional biases. Before the error clamp block, participants were informed about the nature of the perturbation, with the instructions emphasizing that the position of the feedback was independent of their hand movement. There were also three trials to demonstrate the invariant nature of the feedback. For each of these three trials, the target appeared at the 90° location (straight ahead), and on successive trials, the experimenter instructed the participant to “Reach straight to the left” (180°), “Reach straight to the right” (0°), and “Reach backwards towards your torso” (270°). The visual feedback appeared at 90° with respect to the target for all three trials. This was followed by the 600-trial error clamp block. The final two blocks were washout blocks, where the participants were told that no visual feedback will be present, and the instructions emphasized that the participant should reach directly to the target.

### Experiment 1 Data Analysis

The primary dependent variable was endpoint hand angle, defined as the angle of the hand relative to the target when movement amplitude reached 8 cm from the start position (i.e. angle between the lines connecting start position to target and start position to hand). A hand angle of 0° corresponds to a reach directly to the target. Analyses were also performed using heading angle at peak radial velocity rather than endpoint hand angle, which yielded essentially the same results; as such, we only report the results of the analyses using endpoint hand angle. To aid visualization, the hand angle values for the groups with counterclockwise rotations were flipped, such that a positive heading angle corresponds to an angle in the direction of expected adaptation (the opposite direction of the rotated, clamped feedback).

Outlier trials were removed from our analysis using a moving mean with a 5-trial window. Trials in which the hand angles deviated by more than 3.5 standard deviations were excluded from further analysis. These outlier trials were assumed to reflect attentional lapses or trials in which the participant attempted to anticipate the target location, and were excluded from further analysis (less than 1% of all trials and between 0 – 2.3% of trials removed for each participant).

Movement cycles consisted of 4 consecutive reaches (1 reach per 4 target locations). The mean heading angle for each cycle was calculated and baseline subtracted to assess adaptation relative to (small) idiosyncratic biases. Baseline was defined as the last 5 cycles of the veridical feedback baseline block (cycles 16 - 20).

We used three primary measures of adaptation: early adaptation, late adaptation, and aftereffect. Early adaptation was operationalized as the average mean hand angle over cycles 3 – 7 of the perturbation block (Kim et al., 2018). Late adaptation was operationalized as the average mean hand angle over the last 10 cycles of the perturbation block (cycles 161 - 170) (Kim et al., 2018; Morehead et al., 2017). The aftereffect was operationalized as the average mean hand angle over the first cycle of the no-feedback washout block. Note that we opted to use these behavioral measures rather than obtain parameter estimates from exponential fits since the latter approach gives considerable weight to the asymptotic phase of performance and is less sensitive to early differences in rate.

These dependent variables were evaluated with a linear model (two-way ANOVA) to ask whether visual uncertainty has an effect on motor adaptation, and how this effect varies with error size. Pairwise unpaired t-tests were performed, and *p* values were Bonferroni corrected to assess group differences.

### Experiment 2

In Experiment 2, the size and direction of the clamped perturbation was varied from trial to trial, allowing us to examine the effect of uncertainty across a wider range of error sizes. A within-subject design was employed (n=24), with each participant tested in two sessions. In one session, the feedback consisted of a cursor (i.e. one dot), and in the other, the feedback was composed of the cloud of dots. The order of the two types of feedback was counterbalanced across participants, with a gap of one to three days between sessions. Each participant was assigned to reach to a single target, chosen from one of three possible locations (45°, 135°, 225°). The same target location was used for both sessions.

There were four blocks in each session for a total of 1465 trials: No feedback baseline (10 trials), veridical feedback baseline (20 trials), clamped feedback perturbation (1425 trials), and no-feedback post-perturbation (10 trials). During the perturbation block, there were 19 possible positions for the center of the feedback: 0°, ± 1.5°, 3.5°, 10°, 18°, 30°, 45°, 60°, 75°, 90°, with the ± indicating that the perturbation could either be clockwise or counterclockwise from the target. There were 75 trials for each condition, with the size of the perturbation selected at random for each trial.

All other aspects of the experiment were the same as in Experiment 1. Participants were informed of the nature of the clamped feedback prior to the perturbation block. The instructions (together with demonstration trials) emphasized that the feedback position was not contingent on their movement. They were told that the position of the feedback would be randomly determined and that they should ignore it.

### Experiment 2 Data Analysis

Hand angles were measured as in Experiment 1, with each value baseline corrected (subtraction of mean hand angle during last five trials of the feedback baseline block). The primary analysis focused on trial-to-trial changes in hand angle, looking at these values as a function of the size and form of the clamped feedback. Since each participant performed both clockwise and counterclockwise error clamps, we collapsed the data over direction for a given perturbation size, providing a more stable estimate of the Δ hand angle for each condition based on 150 trials/perturbation size, except for the 0° condition which had only 75 trials. Trials in which the change in hand angle that deviated by more than 3.5 standard deviations were excluded from further analyses (less than 1% of all trials and between 0 – 0.8% of trials removed for each participant).

The Δ hand angle data for each of the 10 rotation sizes were submitted to 10 planned paired t-tests, comparing trial-by-trial adaptation between the cursor and cloud conditions. Since the main goal of this study is to identify error sizes where visual uncertainty has an effect on adaptation, applying a Bonferroni family wise error correction on 10 planned comparisons would lead to a loss in power. We therefore corrected for multiple comparison using the less stringent Benjamini-Hochberg Procedure with a false discovery rate of 0.05 (R package: FSA).

### Visual Discrimination Task

To quantify the effect of our uncertainty manipulation, a subset of the participants in Experiment 1 (n = 64) were tested on a perceptual task, comparing position acuity for displays containing a single dot or a cloud of dots (Fig. 2c). For these participants, the visual discrimination task was performed prior to the reaching task.

We used a two-alternative, forced choice visual discrimination task. Each trial began with the presentation of an arrow at the center of the screen that pointed towards one of two possible target locations (45° or 135°) for 1000 ms. Once the arrow disappeared, a blue dot (0.6 mm diameter) was immediately presented at the cued location. This defined the reference position. The referent remained visible for 250 ms, followed by a blank screen for 750 ms. The comparison stimulus was then presented for 500 ms. Using a within-subject design, there were 20 comparison values: 10 displacement sizes (± 0.3°, 0.8°, 1.5°, 2.5°, or 5°) x 2 forms (cursor or cloud, using the same specifications for each as in the reaching phase of the study). Following the offset of the comparison stimulus, the participant vocally indicated if the center of the comparison stimulus was shifted clockwise or counterclockwise relative to the target location. The experimenter entered the participant’s choice with a key press, concluding the trial. A right arrow response was used for clockwise choices and left arrow for counterclockwise choices.

To maintain a similar task context between the visual discrimination task and the reaching task, participants were not asked to maintain fixation. However, we acknowledge that the attentional demands in the two tasks were different. In this visual discrimination task, participants attended to the comparison stimulus to provide an accurate directional judgement relative to the referent, whereas in the reaching task, participants ignored the visual feedback (per the task instructions).

### Visual Discrimination Task Data Analysis

To examine how visual uncertainty influences the assessment of feedback location, we fitted psychometric functions to the participants’ verbal reports. The psychometric function was defined as the probability of reporting “counterclockwise” for each displacement size, *x*. We fit the judgment data with a cumulative density function (cdf) of a normal distribution expressed as *φ*(*x*):

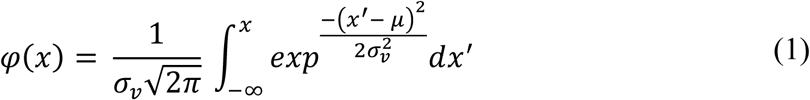

From this function, we obtained the point of subjective equality, *μ*, the mean of the underlying gaussian distribution, as an estimate of the participant’s bias. As a measure of visual uncertainty, we used the difference threshold *σ*_*v*_, the standard deviation of the underlying gaussian distribution. A larger difference threshold is indicative of more variance (and uncertainty) in the perceptual judgments. Pairwise paired t-tests were performed on these two variables to assess within-participant differences in directional bias and visual uncertainty for the cursor and cloud conditions.

### Measures of Effect Size

Cohen’s *d* (for between-subjects design), Cohen’s *d*_*z*_ (for within-subjects design), and eta squared (for between-subjects ANOVA) were provided as standardized measures of effect size (Lakens, 2013). To evaluate key null effects, we calculated the Bayes factor (*BF*_0_+) for the t-values.

## MODELING

In this section, we formalize the four models described in the Introduction. We will express each model in terms of a motor update, the trial-to-trial change in hand angle as a function of the error size.

### The Optimal Integration Model (OI model)

According to the OI model (Burge et al., 2008; Körding & Wolpert, 2004; Wei & Körding, 2010), an estimate of the visuomotor mapping is the integrated effect of two sources of sensory information: visual feedback (*v*) and proprioceptive feedback (*p*). Based on the difference between this estimate and the target location, the amount of trial-to-trial motor update can be computed (*U*). To account for the inherent noise in each sensory modality (*σ*_*v*_ and *σ*_*p*_), the visual and proprioceptive signals are optimally integrated in a maximum-likelihood fashion:

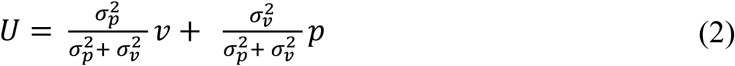

The proprioceptive estimates are assumed to be largely unbiased at the target location during implicit adaptation (i.e., *p* = 0), and the visual feedback on trial *n* is set to the size of the clamped perturbation (i.e., *v* = *e*_*n*_). The expression can be simplified to:

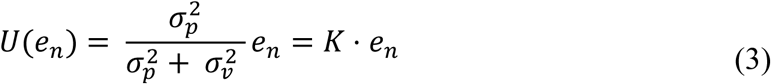

The 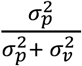 term is known as the Kalman Gain (*K*), which modulates the strength of the error signal and represents the learning rate. Thus, as visual uncertainty increases (*σ*_*v*_ increases), the slope of this motor update function (*K*) and consequent motor updates (*U*) both decrease. The OI model has three free parameters (cursor *σ*_*v*_, cloud *σ*_*v*_, and *σ*_*p*_) that produce a monotonic error-size dependent motor update function (Fig. 1a).

### The Relevance Estimation Model (RE model)

While the OI model assumes that all errors are relevant motor errors, the RE model posits that the system will be more responsive to errors attributed to the agent (e.g., motor execution) and not as responsive to errors attributed to extrinsic factors, at least when the latter tend to be infrequent events. To solve this credit assignment problem, the system performs causal inference, with one heuristic being error size such that the large errors are more likely to be attributed to an extrinsic source.

In this formulation, the motor system updates an internal model of the perturbation by the optimal integration of feedback information (*σ*_*v*_ and *σ*_*p*_) as in the OI derivation, as well as by estimating whether the feedback is relevant, *prob*(*relevant* | *e*_*n*_):

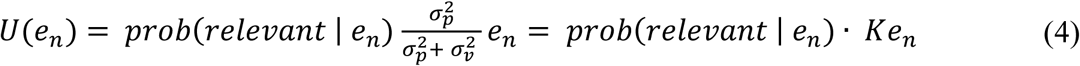

The estimate of error relevancy, *prob*(*relevant* | *e*_*n*_), depends on the size of the error signal, broadened by the noise in each sensory modality (*σ*_*M*_), and scaled by the constant *C* and constant *S*. Thus, following the derivation of Wei and Kording, 2009:

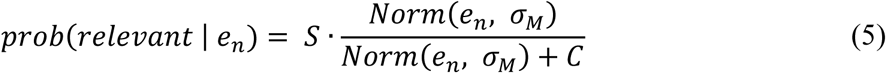

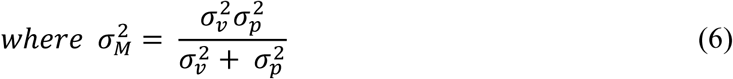

In summary, visual uncertainty (*σ*_*v*_) has two effects on the size of the motor update. Similar to the OI model, visual uncertainty weakens motor updating via a decreased Kalman gain (*K*). But unlike the OI model, visual uncertainty adds uncertainty to causal inference performed by the motor system, making it difficult to definitively attribute errors either to motor errors or extrinsic factors. As a result, visual uncertainty broadens the error relevancy function, *prob*(*relevant* | *e*_*n*_), with the effect that more large errors are attributed to motor errors. Consequently, the RE model predicts a “cross-over” point: While small certain errors induce greater learning than small uncertain errors, large certain errors induce less learning than large uncertain errors (Fig. 1b). The RE model has five free parameters (cursor *σ*_*v*_, cloud *σ*_*v*_, *σ*_*p*_, *S*, and *C*) that produce an error-size dependent non-monotonic motor update function (Fig. 1b).

### The Motor Correction Model

We describe two variants of the MC model (iMC and mMC). Both build on the hypothesis that the motor update function is size-dependent only over a limited range of error sizes and then reaches an invariant, saturated value (*U*_*max*_) for larger errors. The saturation point (*e*_*sat*_) has been estimated to be around 5° (Kim et al., 2018). Given these assumptions, the motor system updates in a bipartite manner:

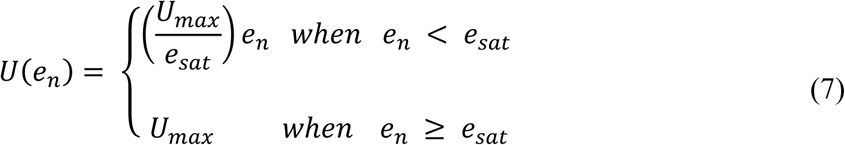

The two variants of the MC model diverge in how visual uncertainty is incorporated. In one variant (iMC model), visual uncertainty is assumed to weaken the motor update for all error sizes, similar to the operation of optimal integration. This can be formalized with a Kalman gain, 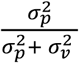, that modulates motor updates based on the relative uncertainty in vision and proprioception, creating an attenuated motor update function, 𝓊_*I*_(*e*_*n*_). The iMC model is a five-parameter model (cursor *σ*_*v*_, cloud *σ*_*v*_, *σ*_*p*_, *e*_*sat*_, *U*_*max*_):

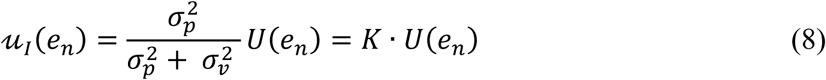

In the second variation, the mis-localization motor correction model (mMC model), visual uncertainty does not change the motor update function. Instead, visual uncertainty adds variability to the perceived location of the feedback, and the update on a given trial will be based on the error size corresponding to the perceived location, rather than the error size corresponding to the actual location (Fig. 1c). We modeled this by sampling the position of the visual feedback from a Gaussian distribution centered at the true error size, *e*_*n*_, with standard deviation, *σ*_*misloc*_, and using the update term associated with that position. The net effect of visual uncertainty over many trials can be calculated by the following expected value of the *U*(*e*_*n*_) function, smoothed over by the Gaussian distribution:

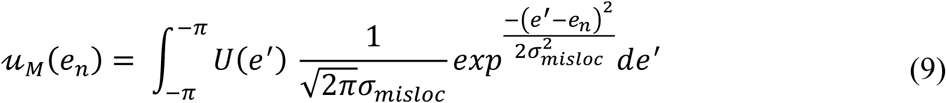

Motor updates to large errors should be essentially insensitive to uncertainty given that the perceived location of the feedback always falls in the saturation zone, and thus the value of 𝓊_*M*_(*e*_*n*_) is the same as the value of *U*(*e*_*n*_). However, for small errors, the update value for the perceived location of the feedback will be different from that associated with the real location. Moreover, the perceived location will sometimes be on the opposite side of the target as the real location, and thus, produce a motor update with the wrong sign. The net effect of these mis-localizations is that learning from small errors will be attenuated. The mMC model is a four-parameter model (cursor *σ*_*misloc*_, cloud *σ*_*misloc*_, *e*_*sat*_, and *U*_*max*_).

### Embedding the Four Models into State Space Equations

As a first test to arbitrate between these four models, we designed experiment 1 to ask how visual uncertainty affects motor adaptation in response to a small (3.5°) or large (30°) error. With a block design, we can quantify the influence of visual uncertainty during the early phase of adaptation and when performance reaches an asymptotic level (Kim et al., 2018). To this end, we embedded all four models within a standard state space equation used to model cumulative learning:

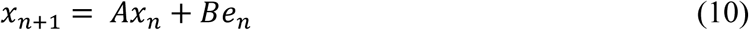

*x* represents the system’s estimate of the perturbation (i.e., and thus produced hand angle) on trial *n. B*, the learning rate, corresponds to the proportion of the error, *e*, used to update the state estimate. With an error clamp, *e* is fixed (e.g., 3.5° or 30° in Experiment 1). *A* represents the retention rate, the amount of learning that is retained from one trial to the next. In all of the model fits, *A* was constrained to be invariant to the form of the error (i.e., same value for the cursor and cloud conditions) and size of the error.

The four models differ in the second half of this equation: the OI model and RE model posit that the learning rate is influenced by feedback uncertainty as formulated by the Kalman gain, *K* (see OI section above):

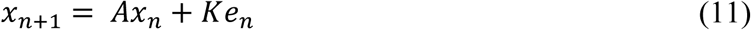

The OI model and RE model differ in their assumption about relevance. The OI model assumes that all sizes of error are relevant (i.e. *prob*(*relevant* | *e*_*n*_) = 1, not explicitly expressed in the equation). In contrast, the RE model assumes that relevance of error depends on the size of the error.

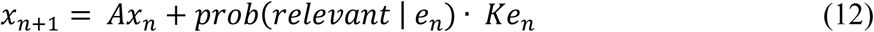

In line with the OI model, the iMC model posits that visual uncertainty, via a decreased Kalman gain, attenuates the entire motor update function:

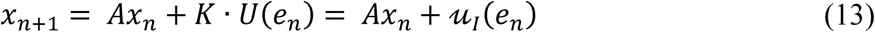

In contrast, the mMC variant posits that visual uncertainty only adds variance to the perceived location of the error:

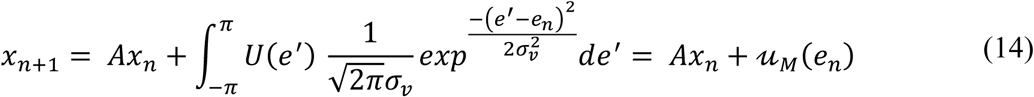

### Model Selection

We used a two-fold cross validation method to evaluate how well each model generalized to independent data sets. We randomly partitioned the data for each condition into two equally disjoint subsets. One subset was used as the training data, creating a model that was tested with the remaining subset. Because the division into folds is random, the two-fold cross-validation procedure was repeated 100 times to provide better estimates of the performance of each model. To quantify model performance, an 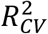 statistic was calculated using the mean of the training data as the null model.

### Parameter Estimation

In order to obtain the best-fitting parameters, we employed a standard jack-knife technique to estimate the mean and confidence interval of the parameter estimates for each model. We first fit all four candidate models to the group averaged data in both experiments to obtain a biased parameter estimate. We then fit the models on *n* group-averaged hand angle data using *n* − 1 samples (*n* is the total number of participants) to estimate the jack-knife estimate of standard errors and bias-corrected jack-knife estimate of each parameter. The specific estimates of each parameter in Exp. 1 were difficult to precisely capture due to co-linearities (i.e., 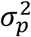 and 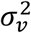 both grow and decay to maintain the same ratio, *K*, in the OI model). Given this, we opted to collapse the right half of the state-space equation (e.g., *Ke*_*n*_) into a single motor update parameter, making it possible to directly compare parameter estimates across the models, and across the two reaching experiments.

## RESULTS

In experiment 1, we asked whether visual uncertainty affects implicit motor adaptation for both small (3.5°) and large (30°) errors. We manipulated uncertainty by presenting endpoint feedback in the form of a cursor (low uncertainty) or cloud of dots (high uncertainty).

### Experiment 1: Perceptual discrimination task

To verify that perceived location is more uncertain with the cloud displays, a subset of participants performed a visual discrimination task before completing the reaching task. For this task, participants compared the relative position of stimuli in two successively presented displays. The first display showed a single dot; the second showed either a single dot or a cloud of dots. The participant judged if the position of the second dot or centroid of the cloud was shifted clockwise or counterclockwise relative to the position of dot in the first display.

As can be seen in Fig. 3a, while participants were mostly correct in assessing the relative positions of both visual stimuli, performance was more variable in the cloud condition compared to the dot condition. To quantify these effects, we estimated the directional bias (*μ*) and the difference threshold (*σ*_*v*_) for each individual in the dot and cloud conditions. Four individuals were excluded from this analysis because their psychometric functions in the cloud condition were too flat to allow the fitting procedure to converge (reflecting poor overall performance).

**Fig. 3.**
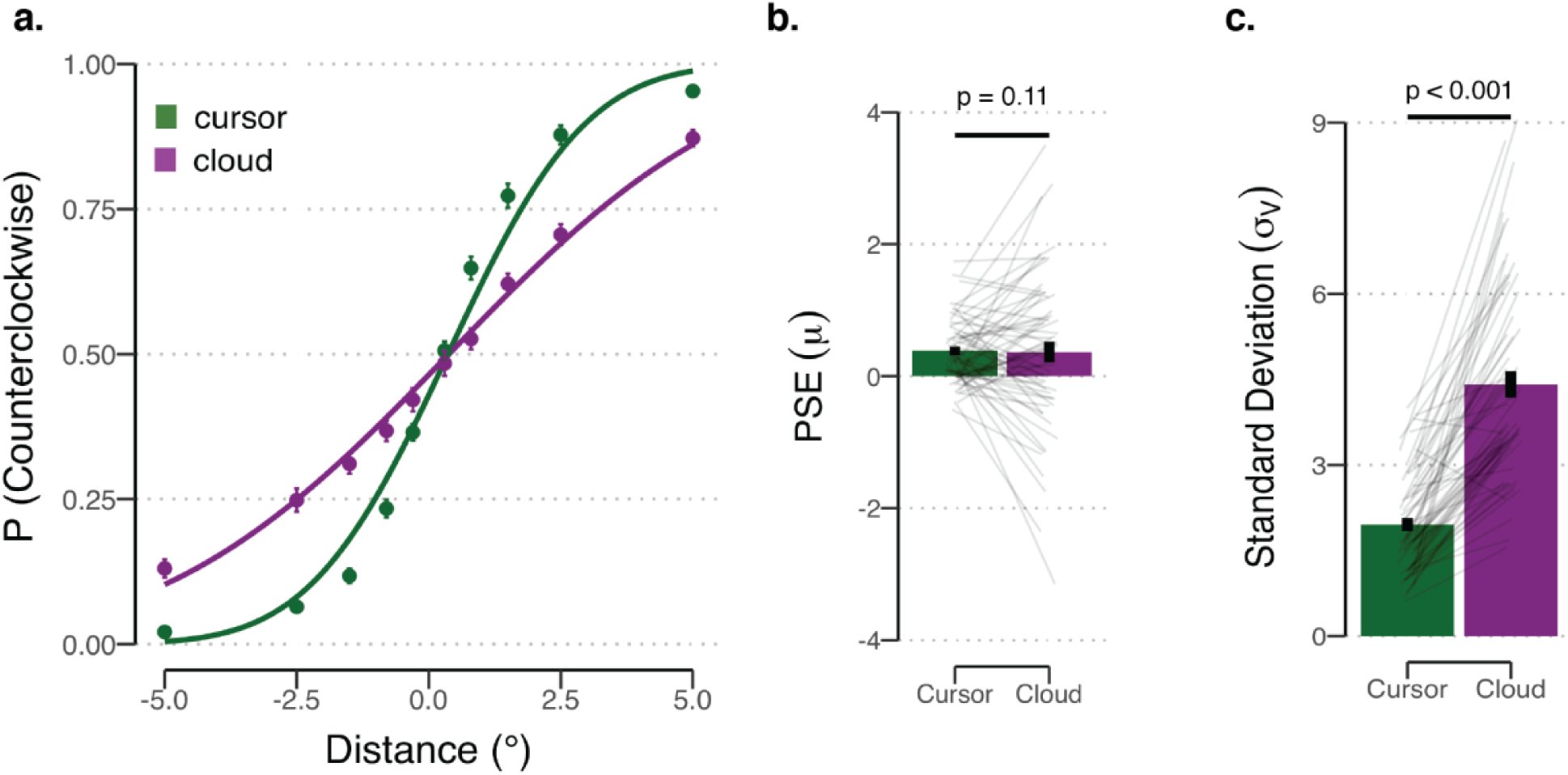
Visual Discrimination Task. **(a)** Proportion of counterclockwise reports as a function of the centroid of the comparison stimulus. Positive values on the x-axis correspond to shifts in the counterclockwise direction. Estimated psychometric functions are the thick lines based on group averaged data for the cursor (green) and cloud (purple) groups. **(b, c)** Mean bias and threshold estimates for the cursor and cloud conditions. Error bars represent SEM and thin gray lines represent individuals’ data.

To test whether there were systematic directional biases, we compared the mean bias estimates to 0° (Fig. 3b). In both conditions, there was a small bias to judge the comparison display as shifted in the counterclockwise direction relative to the referent display (cursor: *t*(58) = 5.71, *p* < 0.001, [0.25, 0.52], *d*_*z*_ = 0.74; cloud: *t*(58) = 2.33, *p* = 0.023, [0.05, 0.68], *d*_*z*_ = 0.30). The degree of bias did not differ between conditions (*t*(58) = 0.13, *p* = 0.90, [−0.27, 0.31], *d*_*z*_ = 0.02).

To test whether the cursor and cloud yield different levels of visual uncertainty, we compared the difference threshold estimates for the two conditions (Fig. 3c). The estimate was considerably higher for the cloud group compared to the cursor group (*t*(58) = −11.66, *p* < 0.001, [−2.88, −2.03], *d*_*z*_ = 1.52). The mean difference thresholds were 1.96° and 4.41° for the cursor and cloud groups, respectively. Thus, the results confirm that participants are more variable in judging the centroid position of a cloud of dots compared to the position of a single dot, a critical assumption underlying our manipulation of visual uncertainty in the reaching experiments.

### Experiment 1: Reaching task

Participants were randomly assigned to one of four groups for the reaching task. After baseline blocks to familiarize the participants with the apparatus and basic trial structure, we presented clamped visual feedback, using either a cursor or a cloud. To assess implicit adaptation, we measured the mean hand angle over the course of the perturbation block and during a subsequent washout block in which no feedback was provided (Fig. 4a).

**Fig. 4.**
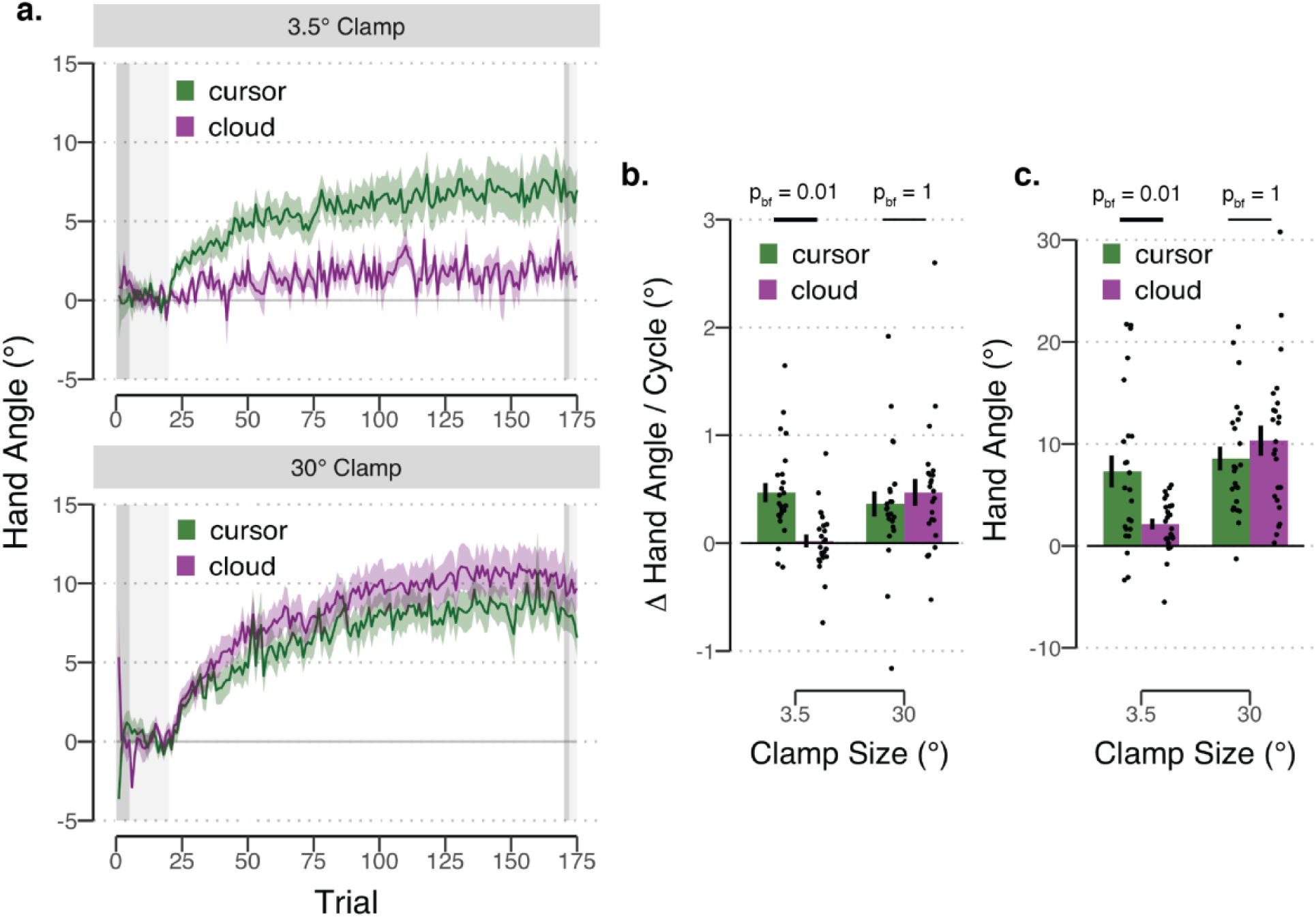
Reaching Results for Experiment 1. **(a)** Mean time course of hand angles for the cursor (green) and cloud (purple) groups, presented with either a 3.5° (top) and 30° (bottom) clamp during the error clamp block. Hand angle is presented relative to the target (light grey horizontal dashed line) during no feedback (dark grey background), veridical feedback (light grey background), and error clamp trials (white background). Shaded region denoting SEM. **(b)** Early adaptation rate, operationalized as the average change in hand angle per cycle over cycles 3 – 7 in the error clamp block. **(c)** Late adaptation, operationalized as average hand angle over the last 10 cycles in the error clamp block.

To test whether the clamped visual feedback elicited implicit adaptation, we first examined the aftereffect results, asking if the mean hand angle was systematically different from 0° in the washout block. The two cursor groups and the 30° cloud group showed implicit adaptation as evidenced by a systematic aftereffect in which the hand angle was shifted in a direction opposite to that of the clamp (3.5° cursor: *t*(22) = 3.98, *p* < 0.001, [3.31, 9.80], *d*_*z*_ = 0.85; 30° cursor: *t*(22) = 7.55, *p* < 0.001, [5.44, 9.66], *d*_*z*_ = 1.5; 30° cloud: *t*(22) = 8.82, *p* < 0.001, [6.43, 11.22], *d*_*z*_ = 1.59). In contrast, the 3.5° cloud group did not show implicit adaptation (*t*(22) = 1.66, *p* = 0.11, [−0.40, 3.72], *d*_*z*_ = 0.34).

We next examined the learning functions, asking how visual uncertainty influenced the rate of early adaptation and the magnitude of late adaptation (Figs. 4b-c). For each dependent variable, we used a 2 × 2 between-subject ANOVA with factors clamp size (3.5° and 30°) and feedback type (cursor and cloud). For early adaptation, neither the effect of clamp size (*F*(1, 92) = 2.93, *p* = 0.09, *η*^2^ = 0.03) nor feedback type (*F*(1, 92) = 2.90, *p* = 0.09, *η*^2^ = 0.03) was significant. However, there was a significant interaction of these factors *F*(1, 92) = 7.53, *p* = 0.007, *η*^2^ = 0.07). Bonferonni corrected *post hoc* analyses revealed that visual uncertainty attenuated adaptation when the clamp size was small (3.5° groups: *t*(46) = 4.16, *p*_*bonf*_ = 0.01, [0.23, 0.67], *d* = 1.20) but not when the clamp size was large (30° groups: *t*(46) = −0.62, *p*_*bonf*_ = 1, [−0.45, 0.24], *d* = 0.17, *BF*_0_+ = 2.97 in favor of the null).

A similar pattern was observed for late adaptation: The clamp size x feedback interaction was significant (*F*(1, 92) = 7.55, *p* = 0.007, *η*^2^ = 0.07), with the post hoc analyses again showing that visual uncertainty attenuated adaptation when the clamp size were small (3.5° groups: *t*(46) = 3.10, *p*_*bonf*_ = 0.03, [1.81, 8.51], *d* = 0.89) but not when the clamp size was large (30° groups: *t*(46) = −0.93, *p*_*bonf*_ = 1, [−5.57, 2.04], *d* = 0.27, *BF*_0_+ = 2.44 in favor of the null). There was also a main effect of clamp size (*F*(1, 92) = 14.04, *p* < 0.001, *η*^2^ = 0.12), with higher asymptotic levels reached for the large clamp compared to the small clamp. The effect of feedback type was not significant (*F*(1, 92) = 1.83, *p* = 0.18, *η*^2^ = 0.02).

We note that overall adaptation in the current study, even in the cursor condition, is attenuated compared to previous studies using the clamp method. Late adaptation to the cursor is between 7 - 9°, values that are much lower than the 20 - 30° asymptotes observed in previous studies using clamped feedback (Kim et al., 2018; Morehead et al., 2017). We suspect the critical difference arises from use of static, endpoint feedback compared to the dynamic, trajectory feedback used in the earlier work. The more ecological nature of an online feedback signal may be a more potent driving signal for learning (Taylor, Krakauer, & Ivry, 2014). Additionally, repeated sampling of the same error signal within a short temporal window may summate into a larger error signal for each movement, resulting in overall higher levels of learning.

In summary, the finding that the effect of visual uncertainty has a large effect on the small clamp size and a negligible effect on the large clamp size poses a challenge to the optimal integration model and the integrated motor correction model. Both of these models predict that increased uncertainty would attenuate adaptation for all clamp sizes, with the effect proportional to the size of the clamp. The absence of an attenuating effect of visual uncertainty for the 30° groups is at odds with this prediction.

### Experiment 2: Reaching task

In the second reaching experiment, we used a within-subject design in which the participants were tested on two separate sessions, once with cursor feedback and once with cloud feedback. Within each session, the error size was varied over a large range (0 to ± 90°). The main dependent variable was the trial-to-trial change in hand angle (Δ hand angle). Positive values indicate a change in hand angle opposite to the direction of the error clamp (i.e., adaptation). Qualitatively, the shape of each function was bipartite, consisting of a roughly linear zone (up to ∼18-30°), followed by an extended saturation zone (Fig. 5).

**Fig. 5.**
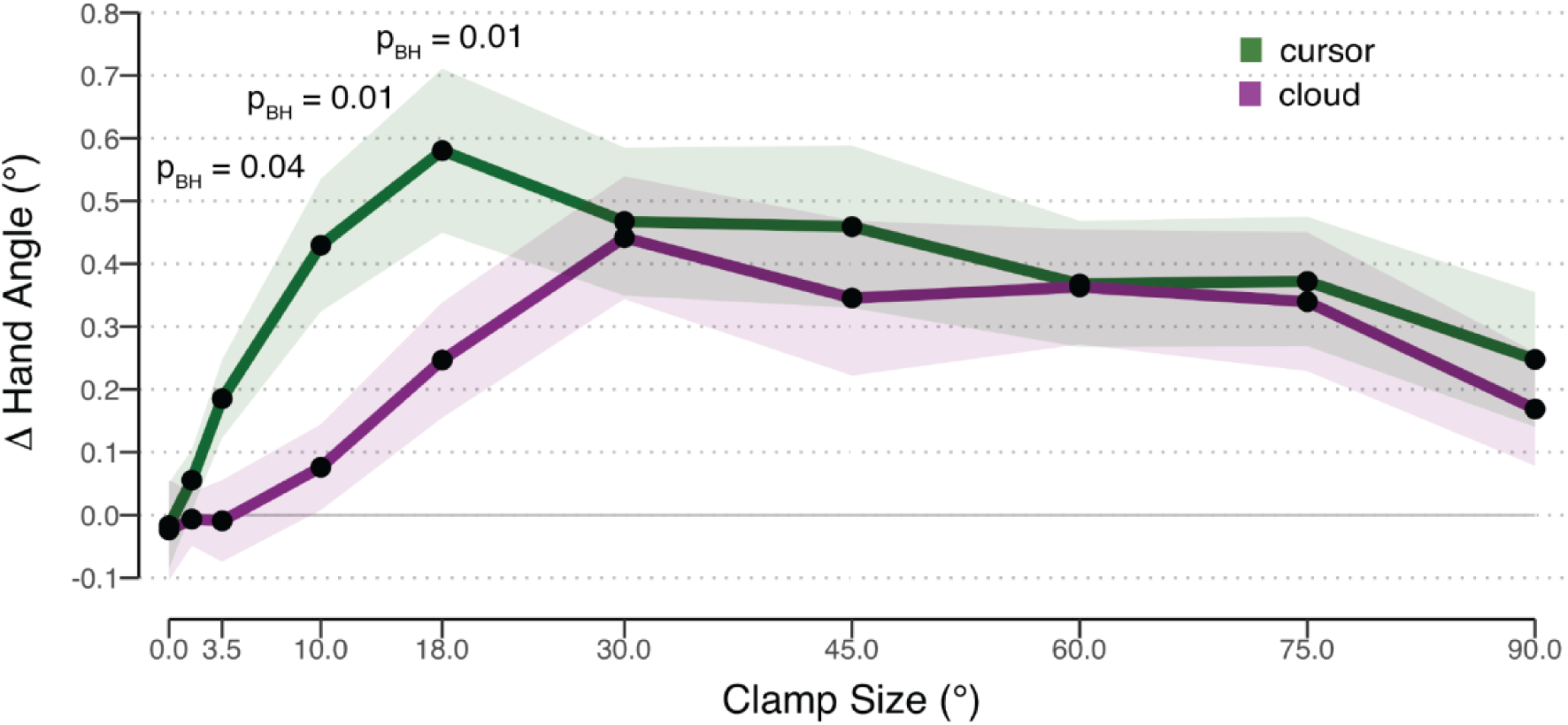
Results for Experiment 2. Change (Δ) in hand angle on trial n + 1 as a function of the clamped feedback on trial n. Feedback was either given in form of a cursor (green) or cloud (purple). Data represent mean corrections across participants. Shaded region represents SEM.

To statistically evaluate these data, we first asked if the Δ hand angle value for each condition (10 error sizes x 2 types of feedback) was significantly different than zero (using a signed test given the prediction that the values would be positive). For the cursor condition (green line), trial-by-trial adaptation was significant for all clamp sizes (all *t*(23) > 2.31, *p* < 0.02, *d* > 0.47) except for the 0° (*t*(23) = −0.25, *p* = 0.60, *d* = 0.05) and 1.5° conditions (*t*(23) = 1.14, *p* = 0.13, *d* = 0.23). In contrast, for the cloud condition, only clamp sizes greater than or equal to 18° elicited significant implicit adaptation (all *t*(23) > 1.86, *p* < 0.04, *d* > 0.38). The trial-to-trial Δ hand angle was not significant when the error was 0° (*t*(23) = −0.30, *p* = 0.62, *d* = 0.06), 1.5° (*t*(23) = −0.15, *p* = 0.56, *d* = 0.03), 3.5° (*t*(23) = −0.15, *p* = 0.58, *d* = 0.03), and 10° (*t*(23) = 1.11, *p* = 0.14, *d* = 0.23).

As can be seen in Fig. 5, the two functions show considerable divergence when the errors are less than 30°, consistent with the observation in experiment 1 that visual uncertainty attenuates adaptation for small clamp sizes. Paired t-tests for each clamp size were used to compare the two feedback conditions. There were no differences in trial-to-trial adaptation in response to clamp sizes greater or equal to 30° (all *t*(23) < 1.29, *p*_*BH*_ > 0.50, *d*_*z*_ < 0.30). In contrast, the cursor elicited a stronger adaptive response than the cloud for clamp sizes of 3.5° (*t*(23) = 2.69, *p*_*BH*_ = 0.04, *d*_*z*_ = 0.55), 10° (*t*(23) = 3.77, *p*_*BH*_ = 0.01, *d*_*z*_ = 0.77), and 18° (*t*(23) = 3.13, *p*_*BH*_ = 0.01, *d*_*z*_ = 0.64) conditions. The mean Δ hand angle was also larger in response to the cursor in the 1.5° condition, but this effect was not significant (*t*(23) = 0.85, *p*_*BH*_ = 0.67, *d*_*z*_ = 0.17).

In summary, the results of experiment 2 provide further evidence that visual uncertainty attenuates adaptation in response to small clamp sizes but has no reliable effect on the response to large clamp sizes. Convergent with the results of experiment 1, this interaction is at odds with the optimal integration and integrated motor correction models, given that these both predict that the attenuated response to high uncertainty feedback will hold for all clamp sizes. Furthermore, the cross-over prediction of the relevance estimation model was also not observed (see Fig. 1b). The results are qualitatively most consistent with the prediction of the mis-localization variant of the motor correction model.

## MODELING RESULTS

We used a cross-validation approach to evaluate the four candidate models described in the Introduction (Fig. 1). For experiment 1, the OI (Fig. 6a) and iMC (Fig. 6c) models qualitatively provided a poor fit to the data. Neither model was able to produce a fit that captured both the attenuated learning for the cloud displays for the 3.5° groups, and similar learning functions for the cloud and cursor groups for the 30° groups. The best fits predicted little to no main effect of feedback type. In contrast, the RE (Fig. 6b) and mMC (Fig. 6d) models were able to capture the interaction. Consistent with these qualitative observations, the OI 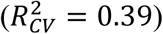 and iMC models 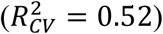 yielded worse fits compared to mMC 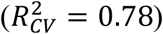 and RE models 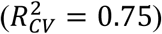.

**Fig. 6.**
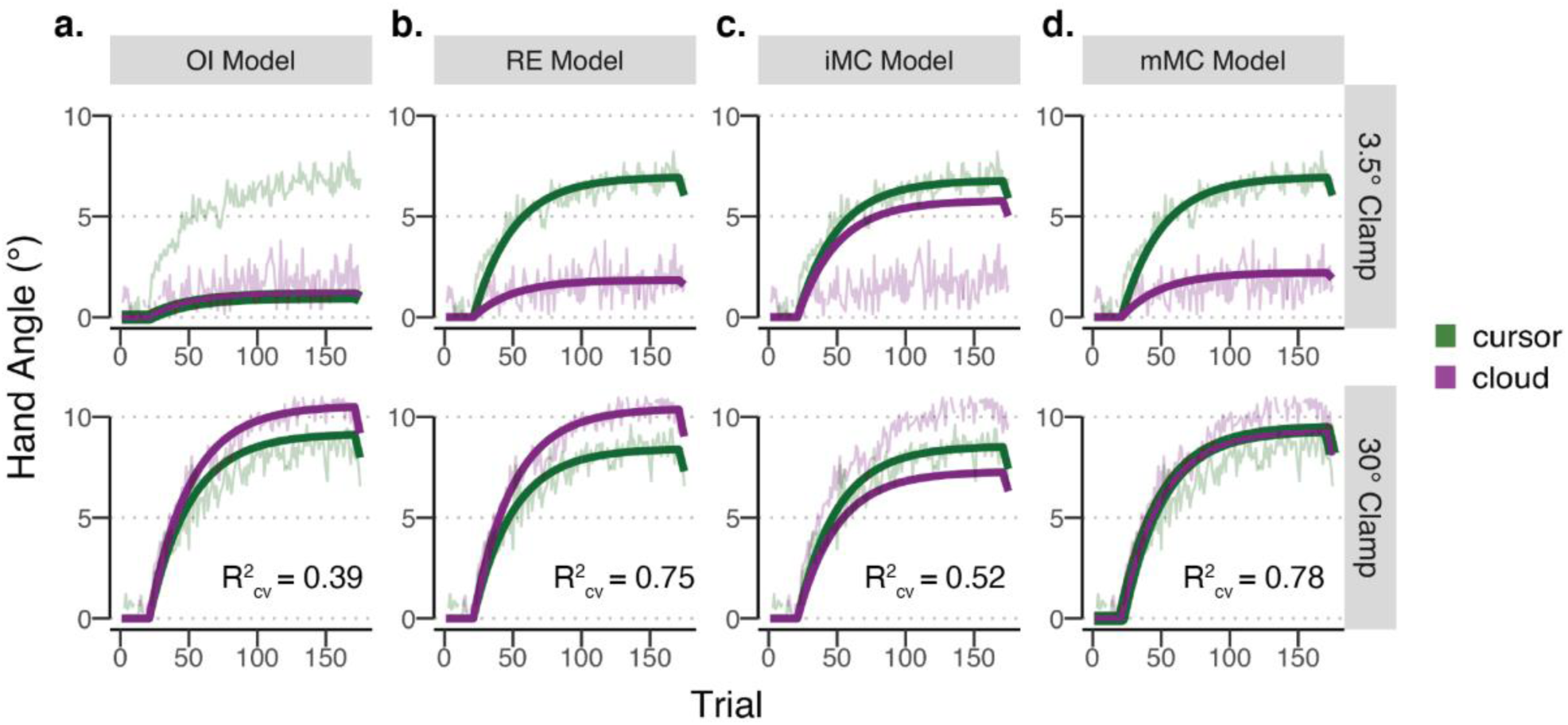
Model Comparison, Experiment 1. Fits of the learning functions from experiment 1, evaluating four hypotheses concerning how visual uncertainty affects adaptation: **(a)** optimal integration model (OI model), **(b)** relevance estimation model (RE model), **(c)** integrated motor correction model (iMC model), and **(d)** mis-localization motor correction model (mMC model). For each model, the plots show the best fit function (solid curves) and group averaged data (lighter, jagged functions), for the cursor (green) and cloud (purple) conditions. Clamp sizes (3.5° or 30°) are delineated by the top and bottom half of the panel, respectively.

The RE and mMC models differ in one important prediction. The mMC model predicts that the learning curve will be the same for the two groups. In contrast, the RE model makes the strong prediction that there will be a cross-over region within which learning will actually be enhanced in response to cloud feedback compared to cursor feedback. This occurs when visual uncertainty makes the motor system broaden its likelihood of relevant errors (i.e., the range of errors attributed to the motor system), and consequently, reduces the system’s discounting of large uncertain errors. Although the mean level of adaptation was greater for the 30° cloud group compared to the 30° cursor group in experiment 1, a tantalizing vote for the RE model, the effect was not significant (*p*_*bf*_ = 1) and largely driven by an outlier in the 30° group (Fig. 4b - c).

To provide a more rigorous test of the cross-over prediction, we sampled a large range of error sizes in experiment 2. The model fits (Fig. 7) were poorer in experiment 2 compared to experiment 1, likely reflecting the noisy nature of trial-by-trial adaptation. Given this caveat, the OI and iMC models again failed to qualitatively capture the interaction of error size and feedback type (OI: 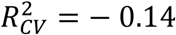; iMC: 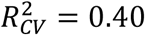), with the best fits here producing a main effect of feedback type (Fig. 7a and 7c, respectively). The best fits were obtained with the RE (Fig. 7b) and the mMC (Fig. 7d) model. Importantly, the fits here yielded a stronger separation of these two models, with the mMC model 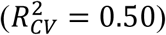 providing a superior fit to the RE model 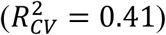.

**Fig. 7.**
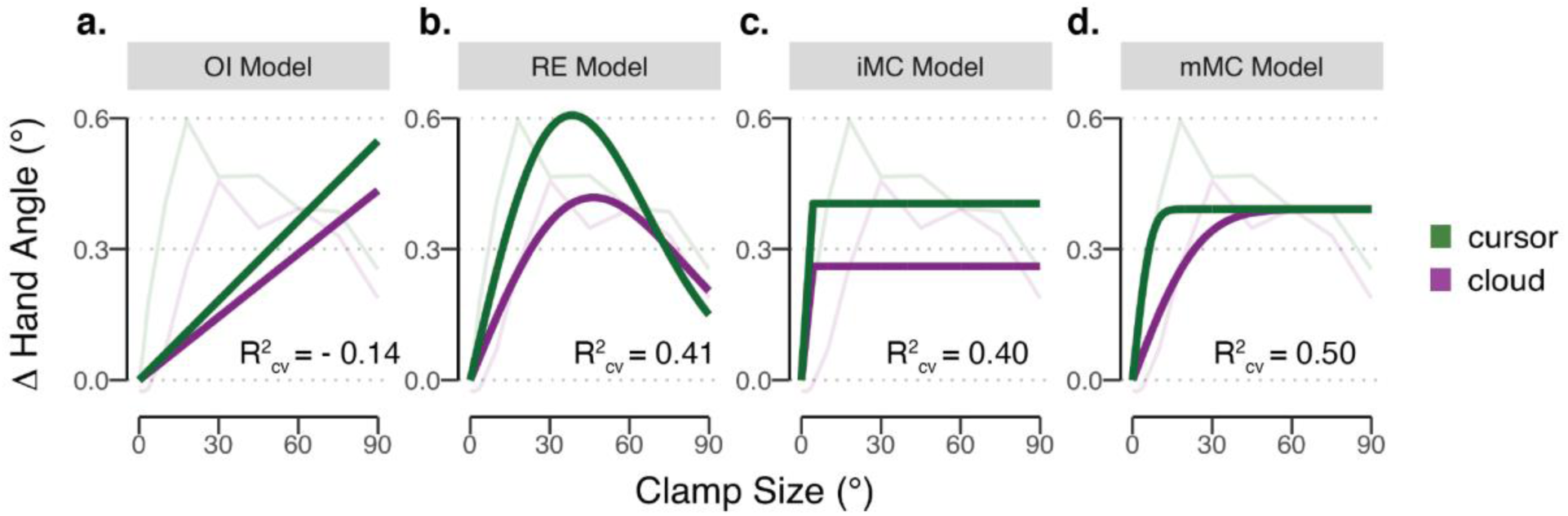
Model Comparison, Experiment 2. Fitted estimates of update size as a function of error size for the four models of adaptation. For each model, the plots show the best fit function (solid curves) and group averaged data (lighter curves), for the cursor (green) and cloud (purple) conditions.

The parameter estimates (Table 1) are also informative in evaluating the four models, and taken as a whole, converge with the fit comparisons in favoring the mMC model over the RE model. One argument is based on parsimony. The mMC model not only provides a better fit but does so with one less parameter than the RE model.

**Table 1.** Model parameters obtained through jack-knife re-sampling techniques for all four models in experiment 2.

| Parameters | OI Model | RE Model | iMC Model | mMC Model |
| --- | --- | --- | --- | --- |
| $\sigma_{v,cursor}$ | **64.17** ± 34.35 | **53.22** ± 8.79 | **10.53** ± 7.53 | |
| $\sigma_{v,cloud}$ | **72.14** ± 38.66 | **85.35** ± 10.36 | **14.68** ± 10.21 | |
| $\sigma_{misloc,cursor}$ | | | | **4.58** ± 1.18 |
| $\sigma_{misloc,cloud}$ | | | | **19.50** ± 4.68 |
| $\sigma_p$ | **5.02** ± 2.71 | **55.38** ± 6.34 | **8.85** ± 5.06 | |
| $U_{max}$ | | | **0.98** ± 0.28 | **0.39** ± 0.10 |
| $e_{saturation}$ | | | **4.40** (*fixed*) | **4.40** (*fixed*) |
| $C$ | | **957.16** ± 316.37 | | |
| $S$ | | **48.17** ± 21.81 | | |

A second argument is based on consistency. The parameter estimates for the mMC model are relatively stable across experiments (Fig. 8). For example, the parameter estimates for motor update across the two experiments follow the same pattern and are of similar magnitude (Fig. 8d). In both, the mean update for the cloud is lower than the cursor in the 3.5° conditions but not when errors are 30°. In contrast, the estimates for the RE model are much more variable, both in terms of the estimated values across the two experiments, but also in terms of the qualitative interaction of the estimated values as a function of clamp size and feedback form. In experiment 1, the estimated update values are similar for the large error and differ for the small error whereas in experiment 2, this pattern reverses.

**Fig. 8.**
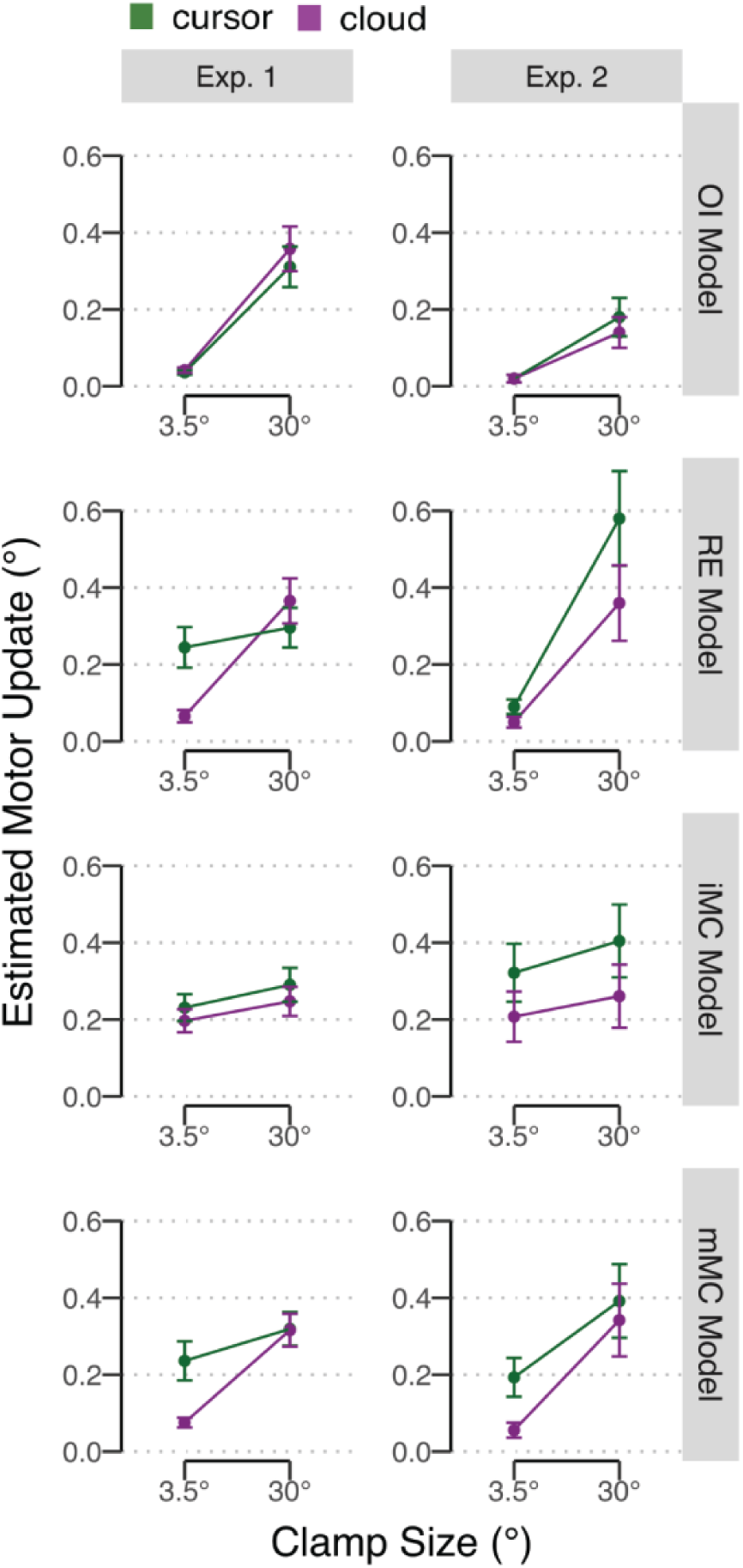
Comparison of Update Size for Comparable Error Sizes in Experiments 1 and 2. Jack-knife estimates of update values in response to 3.5° and 30° cursor (green) or cloud (purple) clamps for the four models. Estimates are based on data from experiments 1 and 2 in the left and right columns, respectively.

A third argument is based on plausibility of parameter estimates. The most direct estimate of the increase in perceptual variability due to the cloud displays comes from the difference thresholds in the perceptual discrimination task. Here the psychometric functions yielded standard deviations of ∼2° for cursor and ∼5° for cloud. For the reaching tasks, the measure of visual uncertainty for each model is obtained by the estimate of variability in determining the centroid of the feedback signal. For the mMC model, the estimates are ∼5° and ∼20° for the cursor and cloud, respectively. Although these values are larger than the perceptual estimations, the attentional state is quite different in the two tasks, with the participants told to focus on the cursor/cloud in the perceptual task and ignore the cursor/cloud in the reaching task. The increase in location variability with a change in attention state is comparable to other reports in the literature (Tsal & Bareket, 1999; Tsal & Bareket, 2005). In contrast, the estimates of visual uncertainty for the RE model are much greater, 53° and 85° for the cursor and cloud, respectively (and in a similar range for the OI model).

## DISCUSSION

Sensorimotor adaptation is affected by the quality of the visual feedback, with the rate of adaptation attenuated as feedback variability increases (Burge et al., 2008; Wei & Körding, 2009, 2010). This effect has been interpreted from an optimal integration framework (Ernst & Banks, 2002): In the context of these visuomotor rotation studies, uncertainty results in less weight being given to the perturbed visual input, relative to that given to veridical proprioceptive feedback, and thus attenuating the resultant change in hand angle. Here we considered an alternative hypothesis, namely that uncertainty does not weaken the strength or relevance of the error signal, but only changes the distribution of the perceived feedback location in an unbiased manner. The results of the two experiments reported here show that visual uncertainty on average attenuated implicit learning in response to small perturbations but had no effect on large perturbations. Complemented by model-based analyses, these results favor the hypothesis that uncertainty increases error mis-localizations rather than modulates error strength.

In the context of optimal integration models (Ernst & Banks, 2002), two fundamental parameters modulate the strength of the error signal in sensorimotor adaptation: the size and the relevance of the error (Burge et al., 2008; Wei & Körding, 2009). However, recent results show that these variables have minimal effects on implicit adaptation (Avraham, Keizman, & Shmuelof, 2019; Morehead et al., 2017). As an alternative, the motor correction model emphasizes constraints on the output of the system, keeping the sensorimotor system exquisitely calibrated to subtle contextual changes (e.g., weight of a worn article of clothing) but within a limited dynamic range. Larger changes require the recruitment of additional learning mechanisms (Bond & Taylor, 2015).

The architecture and operation of the motor correction model requires minimal modification to account for the effect of uncertainty. The system operates in a modular, obligatory manner, taking an estimate of the location of the error to dictate the size of the update. Critically, the system does not respond to distributional information, but functionally operates to convert uncertain error information into a single motor update. This process could proceed in two ways. An estimate of the perceived location is obtained and transformed into a singular motor update. Alternatively, each element of the uncertain feedback (e.g., the dots constituting the cloud) could be associated with its unique update value and these are combined to yield an integrated update value (Kasuga, Hirashima, & Nozaki, 2013). Our experiments cannot distinguish between these alternatives, but they provide interesting hypotheses for future studies.

In both interpretations of the mMC model, the motor system’s response to large errors is unaffected by uncertainty: Even when the distribution of point estimates is large, the associated update values all fall within the saturation zone of the update function. A different picture emerges for small errors, where three types of mis-localizations together attenuate the motor response to noisy feedback. First, some trials will result in an increase in perceived error size, but the associated update values will be constrained by the saturated zone. Second, some trials will result in a decrease in perceived error size, and result in a reduced update size, falling at lower values along the linear zone. Third, on some trials, the perceived error will have the opposite sign, producing an update in the opposite direction.

Taken together, the motor correction model provides a parsimonious account for the effects of visual uncertainty on implicit adaptation. Uncertainty, at least with respect to visual feedback, alters the input to the system, but only in the sense that it alters the range of the perceived location of the feedback. Given that percept, the operation of the update process remains unchanged. Importantly, relevance does not factor in the operation of this learning process, a point emphasized by the striking efficacy of visual error clamps in driving adaptation. The insensitivity of the system to the statistics of the feedback is especially compelling in experiment 1. Here we also attribute the attenuated response to the cloud feedback as resulting, in large part, from trials in which the error was perceived on the opposite side of the target as the actual error. These sign errors occur despite the fact that the actual centroid of the feedback with respect to the target location was invariant for 600 trials.

We note that the mMC model does not consider the contribution of proprioception. The model assumes that the error signal driving learning is dictated solely by the perceived location of the visual feedback. This approximation is reasonable given that visual errors are thought to dominate in the context of a visual perturbation (Sober & Sabes, 2003). However, building on the cue integration idea, *U*_*max*_ could correspond to the point of stability between visual and proprioceptive feedback, rather than the maximum plasticity of the system. Extending the model to incorporate proprioception, including how proprioceptive uncertainty affects learning, is a topic for future studies.

## Author Contributions

All authors contributed to the study design. Testing, data collection, and data analysis were performed by J.S.T. All authors contributed to the interpretation of results under the supervision of R.B.I. J.S.T drafted the manuscript. G.A, H.E.K, D.E.P, and R.B. I. provided critical revisions. All authors approved the final version of the manuscript for submission.

## Acknowledgements

We thank Alan Lee, Cindy Lin, and Noah Bussell for their assistance with data collection. J.S.T was funded by a 2018 Florence P. Kendall Scholarship from the Foundation for Physical Therapy Research. This work was supported by grant NS092079 from the National Institutes of Health.

